# 3’ to 5’ translation of linear mRNAs?

**DOI:** 10.64898/2025.12.29.696803

**Authors:** Xiangyou Pi, Peng Wang, Bin Lin, Tianliang Liu, Zhe Lin, Yanchao Yang, Xiaoru Zhang, Zida Lin, Ling Sun, Fei Chen, Xi Zhang, Zhiliang Ji, Yufeng Yang

## Abstract

3’→5’ translation (termed “backward translation”) mediated by eukaryotic circular RNAs has been reported recently. Whether linear mRNA can produce proteins through backward translation remained elusive. Here, we demonstrate that linear mRNAs can mediate backward translation *in vitro*. Backward translation was detected in human cells transfected with expression vectors used to produce linear mRNAs. Importantly, synthetic linear mRNAs could produce backward translation signal in multiple *in vitro* translation systems, with a level comparable to that of conventional 5’ to 3’ forward translation. Furthermore, we demonstrated that a single translation initiating sequence (TIS) was able to drive backward and forward translation in an *in vitro* experimental setup, with the efficiency of backward translation found to be significant lower than that of forward translation. In summary, this study further reinforces the notion of translation flexibility, demonstrates a broader applicability of backward translation and heralds a new strategy in synthetic biology.

## Introduction

Previously, the groundbreaking finding of 3’→5’ translation (“backward translation”, BT) of eukaryotic circular RNAs (circRNAs) has been reported ^[1]^. In this BT mode, rather than the conventional 5’→3’ translation (i.e. “forward translation”, FT), the triplet codons of circRNAs are decoded along the 3’→5’ end direction to synthesize proteins. The starting 5’-GUA-3’ codon, rather than the conventional 5’-AUG-3’, initiates translation, and translation proceeds via decoding the triplet genetic codes in the 3’→5’ direction along the circRNA till the stop codons 5’-AAU/AGU/GAU-3’ (rather than the conventional 5’-UAG/UGA/UAA-3’) (**Fig. 1**) ^[1]^. Non-Watson-Crick pairing ^[2,3]^ might be the underneath molecular mechanism at the tRNA-RNA interface for the backward translation. This novel translation mechanism has helped disclose unknown proteomes, specifically, nearly 60,000 new proteins/polypetides of high confidence have been revealed for human beings^[1]^. Unique *cis-sequence* features such as backward translation initiator sequences (BTIS) within endogenous circular RNAs have been identified and successfully applied in synthetic biology using circular RNAs as translation templates ^[1]^. However, their relatively low translation initiation efficiency of synthetic backward translatable circular RNAs remains a challenging limitation, possibly due to current limited information on various *cis*- and *trans*-acting regulatory factors. In addition, one key question remains, whether linear mRNAs may might undergo or result in backward translation, which is the focus of our current study.

**Figure 1.**
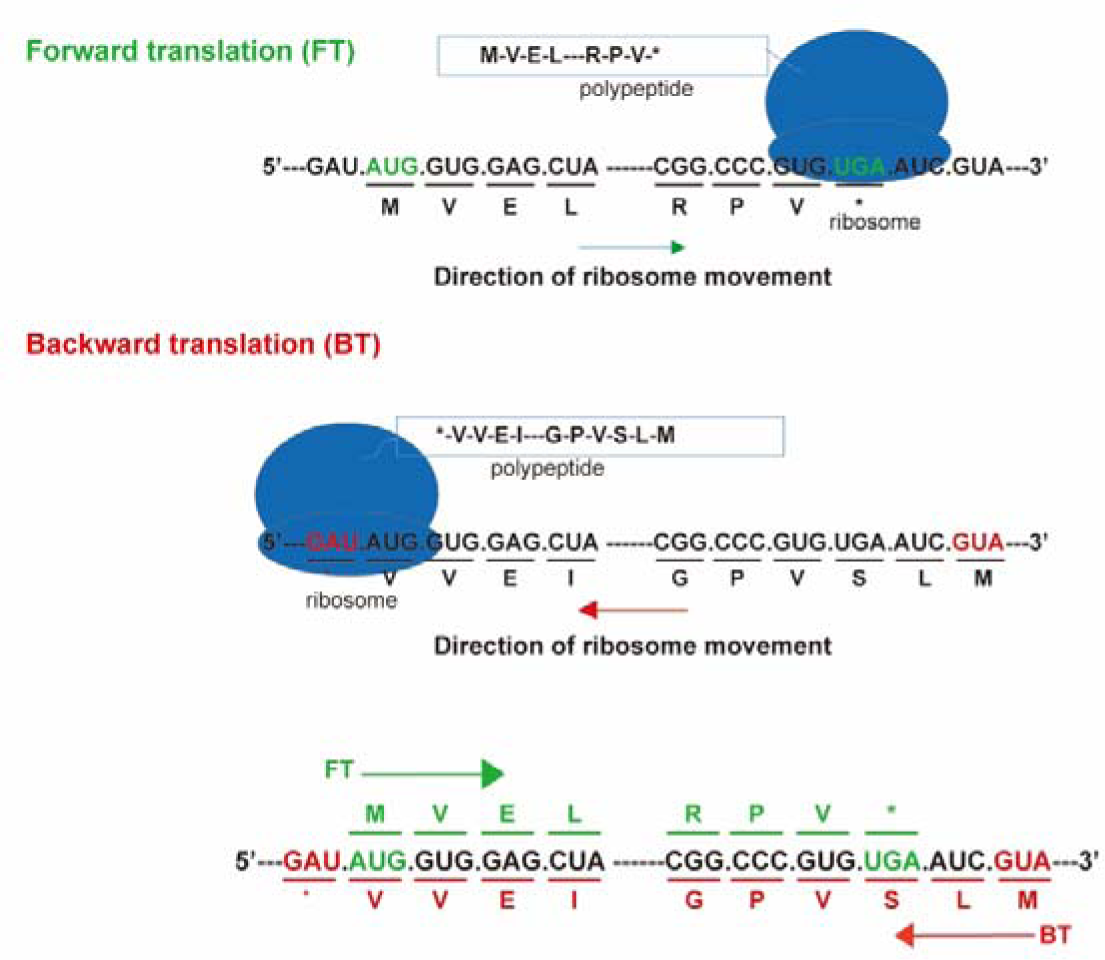
Diagram of 3’ to 5’ backward translation versus conventional 5’ to 3’ forward translation. In this BT mode, rather than the conventional 5’→3’ FT mode, the triplet codons of circRNAs are decoded along the 3’→5’ end direction to synthesize proteins. The starting 5’-GUA-3’ codon, rather than the conventional 5’-AUG-3’, initiates translation, and translation proceeds via decoding the triplet genetic codes in the 3’→5’ direction along the RNA till the stop codons 5’-AAU/AGU/GAU-3’ (rather than the conventional 5’-UAG/UGA/UAA-3’). A virtual RNA template is used for demonstration.

## Results

### Backward translation of linear and circular RNAs resulting from plasmid transfection in HEK-293T cells

We started to repeat the previous results in HEK-293T cells with transfection of DNA vectors that were supposed to be transcribed into backward translatable circRNA. We obtained consistent results, that is, backward translation of circRNAs could be achieved by engineered expression vectors (**Fig. 2A, D**). We then examined whether the same set of TISs ^[1]^, derived from human *hs.circIFITM1*, *hs.circCAPN15* and *Drosophila dm.circSCRIB* respectively, while placed 3’ downstream of backward ORF of eGFP in linear mRNAs could lead to the backward translation of (bw)-eGFP. Excitingly, eGFP-positive cells were detected in all the transfection groups of *hs*.circCAPN15-TIS-(bw)-eGFP, *hs*.circIFITM1-TIS-(bw)-eGFP, and *dm*.circSCRIB-TIS-(bw)-eGFP (**Fig. 2D, E**). These experiments thus provided the first clue for the occurrence of backward translation for linear mRNAs driven by specific *cis*-acting elements.

**Figure 2.**
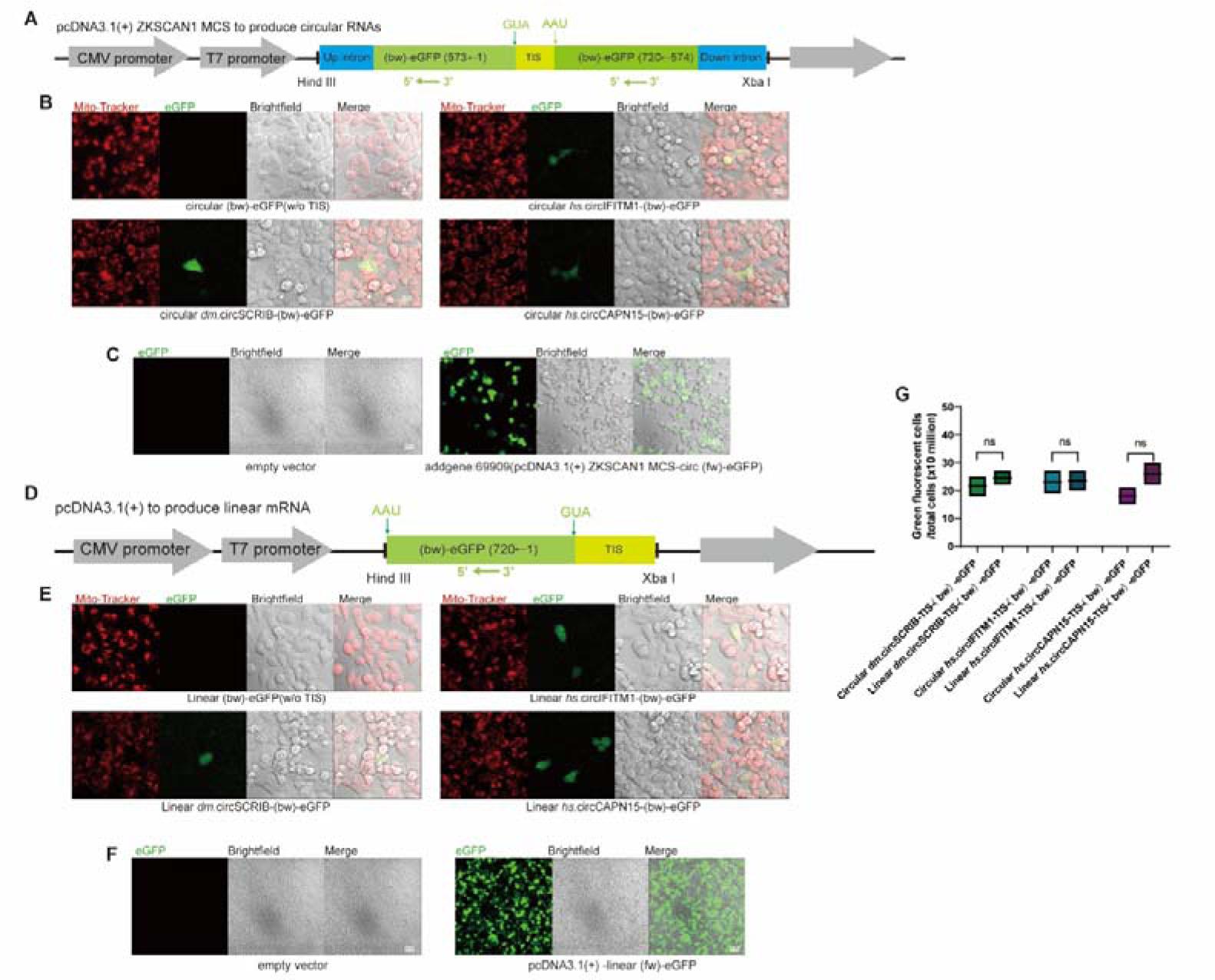
Backward translation of linear and circular RNAs resulting from plasmids transfection in HEK-293T cells. **(A)** Diagram showing pcDNA3.1(+) ZKSCAN1 MCS as the vector for circular mRNA backward translation. The construct consisted of two backward eGFP fragments (coordinates: 593→1 and 720→594), separated by a TIS-containing sequence. The TIS was flanked by a backward start codon (←GUA) and a backward stop codon (AAU←). **(B)** Representative fluorescence microscopy images illustrating positive cells after transfection of HEK-293T cells with circular (bw)-eGFP (w/o TIS), *hs*.circIFITM1-TIS-(bw)-eGFP, *hs*.circCAPN15-TIS-(bw)-eGFP, and *hs*.circSCRIB-TIS-(bw)-eGFP. Scale bar, 20μm. Live cells were stained with Mito-Tracker (red, mitochondria). **(C)** Representative fluorescence microscopy images illustrating HEK-293T cells transfected with the empty vector plasmid, as the negative control. Scale bar, 60μm. HEK-293T cells transfected with the Addgene plasmid #69909 (pcDNA3.1(+) ZKSCAN1 MCS-circ (fw)-eGFP) served as the positive control, with approximately 35%–40% of cells displayed green fluorescence. Scale bar, 40μm. **(D)** Diagram showing pcDNA3.1(+) as the vector for linear mRNA backward translation. A backward GUA (←GUA) and a backward stop codon (AAU←) were inserted before the TIS. **(E)** Representative fluorescence microscopy images illustrating positive cells after transfection of HEK-293T cells with linear (bw)-eGFP (w/o TIS), *hs*.circIFITM1-TIS-(bw)-eGFP, *hs*.circCAPN15-TIS-(bw)-eGFP, and *hs*.circSCRIB-TIS-(bw)-eGFP. Scale bar, 20μm. Live cells were stained with Mito-Tracker (red, mitochondria). **(F)** Representative fluorescence microscopy images illustrating HEK-293T cells transfected with the empty vector plasmid, as the negative control. Scale bar, 60μm. HEK-293T cells transfected with the pcDNA3.1-linear (fw) -eGFP plasmid as the positive control with approximately 85–90% of cells displayed green fluorescence. Scale bar, 60μm. **(G)** Comparison of linear and circular backward translation effiency in the HEK-293T transfection experiments. The Y axis shows the ratio of positive cells after transfection of HEK-293T cells with circular RNA and linear RNA, respectively. Statistical comparisons between groups were performed using Mann-Whitney U in GraphPad Prism (version 9.2.0). Statistical significance: ns: not significant, “*”: p <0.05, “**”: p <0.01, “***”: p <0.001, “****”: p <0.0001.

To further evaluate the efficiency of this process, we compared the backward translation efficiency these TISs in the linear mRNA contexts with that of circular RNA contexts. RT-PCR analysis confirmed the expression of these engineered linear mRNAs and circular RNAs in transfected cells (**Fig. S1**). We found that the backward translation efficiency in linear mRNAs was comparable to that in circular RNAs, albeit both occurred at extremely low levels (**Fig. 2G**). This finding suggests that the occurrence of backward translation might depend primarily on the core function of the TISs sequence itself and might not be sensitive to the origin topological configuration (linear or circular) of the RNA template.

### *In vitro* backward translation mediated by synthetic linear mRNA templates

Furthermore, to rule out potential false-positive arising from aberrant plasmid expression, we synthesized *hs*.circIFITM1-TIS-(bw)-Gaussia and *dm*.circSCRIB-TIS-(bw)-Gaussia linear mRNAs via T7 *in vitro* transcription (**Fig. 3A**). We then examined whether synthetic linear mRNAs could result in backward translation using the commonly employed *in vitro* translation system, the cell-free rabbit reticulocyte lysate (RRL) (**Fig. S2A**). Compared to the negative control (no mRNA), the luminescence intensity increased by 22.33% for *dm*.circSCRIB-TIS-(bw)-Gaussia and by 23.98% for *hs*.circIFITM1-TIS-(bw)-Gaussia, while no signal was detected for (bw)-Gaussia without a TIS, indicating TIS-dependent linear mRNA backward translation activity (**Fig. 3B**). To verify that linear backward translation is not limited to RRL, we further tested the activity of linear *dm*.circSCRIB-TIS-(bw)-Gaussia using cell-free *Drosophila* ovary extract^[4,5]^. We observed an approximately two-fold increase in luminescence intensity compared to that in the negative control (no mRNA) (**Fig. 3C**). The signal strength from linear mRNAs designed for backward translation was comparable to that of conventional forward translation mediated by m7G-CrPV-IRES-Gaussia, yet slightly weaker (**Fig. 3C**). Furthermore, using a reconstituted ribosome complex ^[6,7]^, we also detected positive signals from synthetic linear *dm*.circSCRIB-TIS-(bw)-Gaussia mRNA (**Fig. 3D and Fig. S2B**).

**Figure 3.**
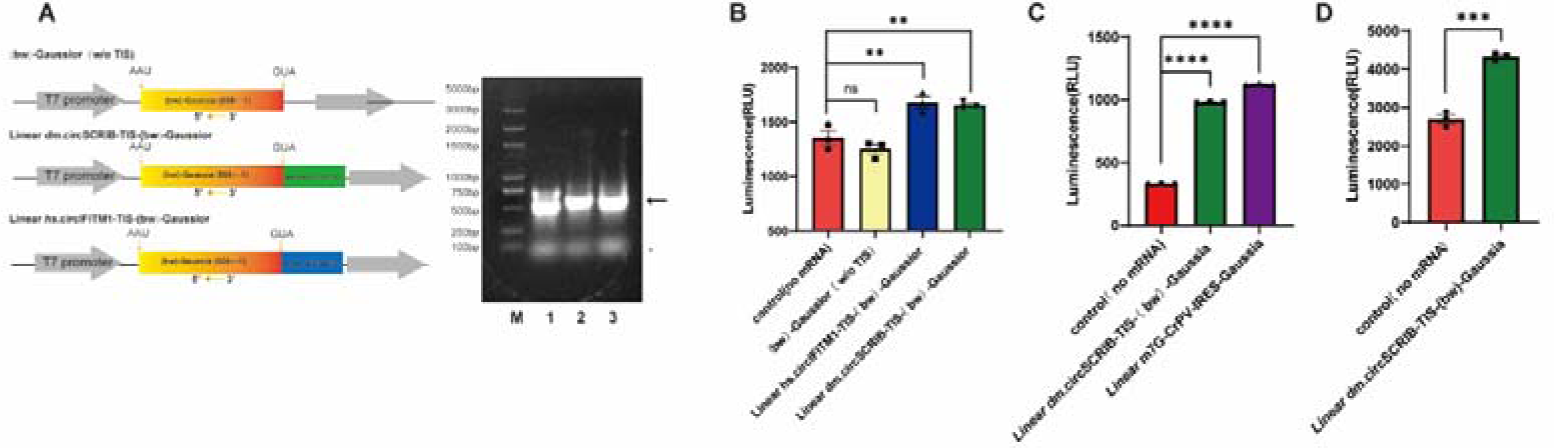
*In vitro* backward translation mediated by synthetic linear mRNA templates. **(A)** Diagram showing the linear mRNA templates for backward translation. The agarose gel electrophoresis on the right shows the mRNA templates after T7 *in vitro* transcription. Lane 1: (bw)-Gaussia (w/o TIS), approximately 720 nt. Lane 2: linear *dm*.circSCRIB-TIS-(bw)-Gaussia, approximately 944 nt. Lane 3: linear *hs*.circIFITM1-TIS-(bw)-Gaussia, approximately 891 nt. A 5000bp ladder marker was used. The asterisk (*) denotes non-specific bands. **(B)** Rabbit reticulocyte lysate was used as the *in vitro* translation system to detect backward translation activity of synthetic linear mRNA templates. Synthetic linear *hs*.circIFITM1-TIS-(bw)-Gaussior and linear *dm*.circSCRIB-TIS-(bw)-Gaussior RNAs were assayed (n=3). Translation was measured by chemiluminescence from Gaussior luciferase activity (coelenterazine substrate, λ =480 nm). Negative control (no mRNA), (bw)-Gaussior (w/o TIS) represents the linear Gaussior RNA without any TIS. **(C)** *Drosophila* ovarian extract system was used as the *in vitro* translation system to detect backward translation activity of synthetic linear mRNA templates. Synthetic backward translatable linear *dm*.circSCRIB TIS-(bw)-Gaussia mRNA (n=3), and synthetic conventional forward translatable linear capped (m7G) Gaussia mRNA containing the Cricket paralysis virus internal ribosome entry site (CrPV IRES) were assayed. Negative control (no mRNA). **(D)** Reconstituted ribosome complex was used as the *in vitro* translation system to detect backward translation activity of synthetic linear *dm*.circSCRIB-TIS-(bw)-Gaussia mRNA. Negative control (no mRNA). Statistical comparisons between groups were performed using Mann-Whitney U in GraphPad Prism (version 9.2.0). Statistical significance: ns: not significant, “*”: p <0.05, “**”: p <0.01, “***”: p <0.001, “****”: p <0.0001.

Taken together, these TIS-dependent reporter assays across multiple cell-free systems provided conclusive *in vitro* biochemical evidence for the occurrence of backward translation mediated by linear mRNA templates.

### A single TIS directs both forward and backward translation

To directly compare the efficiencies of canonical forward translation and backward translation, we constructed a dual-fluorescence reporter system along a single linear mRNA template. In this system, the open reading frames (ORFs) for a forward red fluorescent protein (mScarlet3) and a backward green fluorescent protein (eGFP) were positioned downstream and upstream, respectively, of a translation initiation sequence (TIS) within a single transcriptional unit (**Fig. 4A**). Quantitative analysis following transfection of HEK-293T cells with reporters containing different TISs revealed the following results: For the *hs*.circIFITM1-TIS-derived construct, 26.8% of cells were positive for forward translation (mScarlet3), while backward translation (eGFP) was extremely low, detected in only 2-3 out of 100,000 cells. Among these rare eGFP-positive cells, 60.0% co-expressed mScarlet3 (**Fig. 4B**); For the *dm*.circSCRIB-TIS construct, 20.0% of cells were mScarlet3-positive, with eGFP-positive cells again at 2-3 per 100,000, and 70.0% of these were dual-positive (**Fig. 4C**). For the *hs*.circCAPN15-TIS-derived construct, ∼30.0% of cells were mScarlet3-positive, 2-3 eGFP positive cells per 100,000, with 60.0% of green cells being dual-positive (**Fig. 4D**). Note that cells with low fluorescence protein abundance and below the detection sensitivity of the microscope were deemed as negative in these experiments, thus, cells with backward translation might be under-estimated.

**Figure 4.**
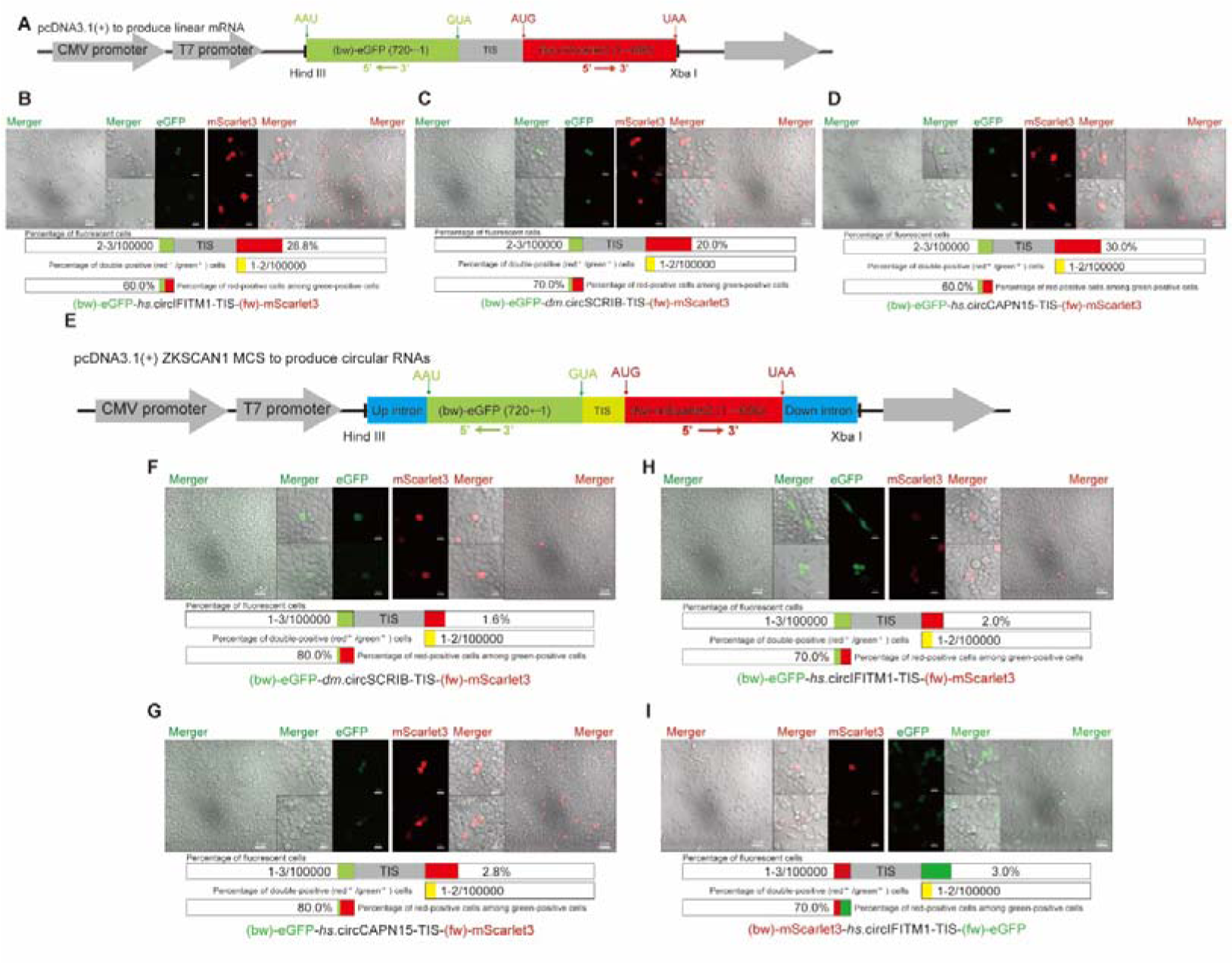
Bidirectional translation mediated by a single TIS. **(A)** Diagram showing design of the pcDNA3.1(+) vector for bidirectional translation of linear mRNAs. The backward (bw)-eGFP was placed upstream of the TIS, and the forward (fw) mScarlet3 was placed downstream of the TIS. **(B)** Representative fluorescence microscopy images illustrating HEK-293T cells transfected with the constructed linear bidirectional fluorescence (bw)-eGFP-*hs*.circIFITM1-TIS-(fw)-mScarlet3. Scale bar, 20 µm. **(C)** Representative fluorescence microscopy images illustrating HEK-293T cells transfected with the constructed linear bidirectional fluorescence (bw)-eGFP-*dm*.circSCRIB-TIS-(fw)-mScarlet3. Scale bar, 20 µm. **(D)** Representative fluorescence microscopy images illustrating HEK-293T cells transfected with the linear constructed bidirectional fluorescence (bw)-eGFP-*hs*.circCAPN15-TIS-(fw)-mScarlet3. Scale bar, 20 µm. **(E)** Diagram showing design of the pcDNA3.1(+) ZKSCAN1 MCS vector for bidirectional translation of circular RNAs. The backward (bw)-eGFP was placed upstream of the TIS, and the forward (fw) mScarlet3 was downstream of the TIS. **(F)** Representative fluorescence microscopy images illustrating HEK-293T cells transfected with the constructed circular bidirectional fluorescent circular (bw)-eGFP-*dm*.circSCRIB-TIS-(fw)-mScarlet3. Scale bar, 20 µm. **(G)** Representative fluorescence microscopy images illustrating HEK-293T cells transfected with the constructed circular bidirectional fluorescent (bw)-eGFP-*hs*.circCAPN15-TIS-(fw)-mScarlet3. Scale bar, 20 µm. **(H)** Representative fluorescence microscopy images illustrating HEK-293T cells transfected with the constructed circular bidirectional fluorescent (bw)-eGFP-*hs*.circIFITM1-TIS-(fw)-mScarlet3. Scale bar, 20 µm. **(I)** Representative fluorescence microscopy images illustrating HEK-293T cells transfected with the constructed circular bidirectional fluorescent (bw)-mScarlet3-*hs*.circIFITM1-TIS-(fw)-eGFP. Scale bar, 20 µm. Quantifications of the fluorescence positive cells ratio after transfection were shown below each microscopy images panel.

We then examined the dual-fluorescence reporter in circular RNAs (**Fig. 4E**). For the circular *dm.*circSCRIB-TIS reporter, forward translation (mScarlet3-positive) was observed in 1.60% of transfected cells, whereas backward translation (eGFP-positive) occurred at 1–3 cells per 100,000. Among these rare eGFP-positive cells, 80.0% co-expressed mScarlet3 (**Fig. 4F**). A similar pattern was observed for the circular *hs*.circCAPN15-TIS reporter: 2.8% of cells were mScarlet3-positive, 1–3 eGFP-positive cells per 100,000, and 80.0% of these were dual-positive (**Fig. 4G**). For the circular *hs*.circIFITM1-TIS reporter, 2.0% of cells were mScarlet3-positive, the eGFP signal remained at a baseline low level (1–3 cells per 100,000), and 70.0% of eGFP-positive cells were also mScarlet3-positive (**Fig. 4H**). To rule out potential sequence- or fluorescence-specific artifacts, we swapped the fluorescent protein positions in the *hs*.circIFITM1-TIS circular reporter, placing a mScarlet3 ORF upstream and a eGFP ORF downstream of the TIS. In this configuration, forward translation (eGFP-positive) was observed in 3.0% of cells, while the signal from backward mScarlet3 remained low (1–3 cells per 100,000), with 70.0% of the mScarlet3-positive cells being dual-positive (**Fig. 4I**), confirming that the observed bidirectional translation asymmetry was independent of the specific fluorescent protein used. Therefore, our experimental data demonstrated that within this particular template, forward translation of linear or circular RNA remarkably outperformed backward translation.

## Discussion

This study established that linear mRNAs, can result in backward translation, indicating that for the translation machinery, there is no fundamental difference between circular and linear RNAs in the process of backward translation. We speculate that backward translation of circular and linear RNAs likely shares a very similar set of translation molecular machinery and regulatory mechanisms. Thus, this current finding demonstrates a broader applicability of backward translation and also indicates that the covalently closed circular structure of circular RNAs might be not a necessary condition for initiating backward translation. On the other hand, it is worth further rigorous investigations whether lariat or other complicated configurations of linear mRNAs or the closed-loop structure formed by the conventional linear mRNA 5’ cap, 3’ polyA and their associated proteins has similarities with that of circular RNAs in terms of backward translation capacity. This also indicates that we can not make an exclusive conclusion that “linear” mRNA directly undergo backward translation at present. However, if we combine the current study on linear mRNAs and the previous report on circular RNAs, the statement of “backward translation of RNAs” sounds more reasonable. In summary, these results further support the notion that ribosome-centered translation machinery has flexibility in translation depending on the RNA template ^[8]^.

Furthermore, we cannot currently completely rule out the possibility that circular RNAs and linear RNAs have some unique regulatory mechanisms for backward translation, including *cis* elements and *trans* factors, which are supposed to be subject to selective pressure during evolution. Our ongoing investigations, such as systematic RNA-binding proteins identification, whole-genome-wide screening, cryo-electron microscopy structural analysis, single molecule imaging study, etc., will help address this interesting question.

Although current study demonstrates that synthetic linear mRNA templates can mediate backward translation *in vitro*, whether endogenous linear mRNA can do so remains elusive. With our initial efforts, BT mediated by eukaryotic endogenous linear mRNAs has not yet been unambiguously validated. At present, we tend to think that naturally occurring linear cases are very rare in eukaryotes. However, in the realm of viruses, due to their compact genomes, backward translation might very likely be utilized, making it an interesting area worth exploring in the near future

Comparable levels of translation signal derived from backward and forward ORFs were detected in the *in vitro* translation system (**Fig. 3C**), on the contrary, several orders of magnitude difference between forward and backward translation was observed in the dual-fluorescence experiments (**Fig. 4**). Due to the limited TISs examined and different assays used in this study, and most importantly, due to the current limited information about the mechanisms of backward translation, we are currently unable to reconcile the seemingly contradictory phenomena observed in the above experiments and reach a general conclusions yet.

Nevertheless, evolutionary pressure may favor gene sequence corresponding to the more energy-efficient forward translation mode, because from a thermodynamic perspective, conventional Watson-Crick pairing is superior to non-Watson-Crick pairing, which could be demonstrated by molecular dynamic simulations. This preference can also explain why endogenous backward translation of linear mRNA may be very rare, making it really fascinating why and how circular RNA backward translation has managed to survive during evolution. Mechanistically, how the translation machinery adapts to the non-Watson-Crick pairing interface of tRNA-mRNA remains to be elucidated, hopefully with the help of cryo-electron microscopy structural analysis study we are now conducting. Meanwhile, systematic identification of *cis* elements and *trans* factors in backward translation mentioned above shall be beneficial for elucidating the regulatory mechanisms, improving the efficiency of *in vitro* translation, which in turn aids in optimizing strategies in synthetic biology.

## Methods and Materials

### Cell culture

HEK-293T cell lines were authenticated by STR analysis. Cells were cultured in DMEM medium (Gibco, C11995500BT), supplemented with 10% fetal bovine serum (Gibco, ST30-3302) and 1% penicillin-streptomycin (Beyotime, 15140-122), and incubated at 37°C in a humidified incubator with 5% CO_2_.

### Plasmid construction and preparation

#### Linear (bw)-eGFP plasmid construction

The mammalian expression vector pcDNA3.1(+) was linearized by double digestion with the restriction enzymes HindIII and XbaI. The backward (bw)-eGFP cassette (5’-AAU← GUA-3’) was inserted upstream of the translation initiator sequences (TIS), resulting in the final construct named pcDNA3.1(+)-(bw)-eGFP-TIS.

#### Linear dual-fluorescence plasmid construction

The mammalian expression vector pcDNA3.1(+) was linearized by double digestion with restriction enzymes HindIII and XbaI. A backward (bw)-eGFP cassette (5’-AAU←GUA-3’) was inserted upstream of the translation initiator sequences (TIS), and a forward (fw)-mScarlet3 cassette (5’-AUG→UAA-3’) was inserted downstream of the TIS. This assembly yielded the final construct named pcDNA3.1(+)-(bw)-eGFP-TIS-(fw)-mScarlet3.

#### Circular (bw)-eGFP plasmid construction

The vector pcDNA3.1(+) ZKSCAN1 MCS + Sense IRES (Addgene: #69909) was linearized using HindIII and XbaI. An Upstream intron (Up-intron) and the backward (bw)-eGFP (573←1) were inserted upstream of the TIS, while the (bw)-eGFP (720←574) and a Downstream intron (Dow-intron) were inserted downstream of the TIS.

#### Circular dual-fluorescence plasmid construction

The vector pcDNA3.1(+) ZKSCAN1 MCS + Sense IRES (Addgene: #69909) was linearized using HindIII and XbaI. An Upstream intron (Up-intron) and the backward (bw)-eGFP cassette (5’-AAU←GUA-3’) were inserted upstream of the TIS, while the forward (fw)-mScarlet3 cassette (5’-AUG→UAA-3’) and a Downstream intron (Dow-intron) were inserted downstream of the TIS. The Up-intron and Dow-intron sequences serve as intronic circularization sequences to facilitate the generation of circular RNA. The resulting construct was named pcDNA3.1(+)-Up-intron-(bw)-eGFP-TIS-(fw)-mScarlet3-Dow-intron.

### Plasmid Extraction and Transfection into HEK-293T Cells

The compositions of plasmids for human cell transfection designed in this study are illustrated in **Fig. 2A, 2D** and **Fig.4A, 4E**. The plasmid was extracted using Endo-Free Midi Plasmid Kit (TIANGEN, #DP108). The plasmids were transfected into HEK-293T cells via an electroporation kit (Celetrix, 1216) according to the experimental needs. For electroporation of HEK-293T cells, the voltage was set to 600V with a duration of 30ms. Six hours post-electroporation, the medium was changed to remove the electroporation solution. The backward translation initiator sequence (TIS) and plasmid primer information used in this study are compiled in **Supplementary Table 1**.

### *In vitro* transcription of RNA

*In vitro* transcription of RNA was performed using the T7 High-Efficiency Transcription Kit (Hzyms, HBP001510) following the manufacturer’s protocol. Briefly, linearized DNA templates were incubated with the provided reaction mix containing T7 RNA polymerase at 37 °C for 2 hours. The Hzymes kit contains cap analogies. The Cap GAG analog has the structure m7G(5’)ppp(5’)(2’OMeA)pG and is suitable for co-transcription capping reactions. It exhibits higher capping efficiency compared to ARCA, enabling the efficient production of natural Cap 1 structured mRNAs. As a result, it helps reduce the immunogenicity of mRNA *in vivo*. Subsequently, the DNA template was removed by digestion with DNase I at 37 °C for 30 minutes. The transcription reaction was terminated by the addition of 3 M sodium acetate (pH 5.2). The total volume was adjusted to 200 µL with nuclease-free water. RNA was then purified by phenol-chloroform extraction followed by precipitation with isopropanol. The final RNA pellet was washed twice with 75% ethanol, air-dried briefly, and resuspended in an appropriate volume of nuclease-free water. The concentration and purity of the synthesized RNA were determined by spectrophotometry.

### RT-qPCR/RT-PCR

Total RNA was extracted from HEK-293T cells using RNAiso Plus (TaKaRa, 9108) according to the manufacturer’s instructions. The total RNA was homogenized thoroughly, and the gDNA was removed using DNase I. For circular RNA, RNase R (Beyotime, R7092S) treatment was performed according to the manufacturer’s instructions to degrade linear RNA. Reverse transcribe clean circRNA into cDNA using a reverse transcription kit with random hexamers (Thermo Scientific, K1622). Design divergent primers for RT qPCR (**Supplementary Table 1**) to quantify circRNAs. For linear RNA, reverse transcribe cDNA directly using a reverse transcription kit and quantify linear RNA using primers. Simultaneously, following conventional PCR amplification with these primers, the products were analyzed and verified by 1.5% agarose gel electrophoresis.

### *In vitro* translation assay

To assess backward translation from linear RNAs, we used the Rabbit Reticulocyte Lysate System (Promega, L4960) for Gaussia protein synthesis mediated by the TIS element. *In vitro* translation was performed with 2.5 μg of RNA incubated at 30°C for 120 minutes according to the manufacturer’s instructions. Luminescence was measured using the Gaussia-Lumi™ Luciferase Reporter Assay Kit (Beyotime, RG072S). For detection, Gaussia-Lumi™ substrate and buffer were mixed at a 1:100 ratio to prepare the working solution. A total of 10μL of translation product was mixed with 100μL of detection reagent (final volume: 110μL), incubated for 5 minutes at room temperature in the dark, and chemiluminescence was recorded using the GloMax Multi Jr detection system (Promega, E6070) with an integration time of 10 seconds per well.

### Ribosome complex reconstitution

Ribosome Isolation from Rabbit Reticulocyte Lysate: A 10%-50% sucrose gradient buffer (SGB; 20 mM Tris-HCl pH 7.5, 150 mM KCl, 5 mM MgCl_2_, 1x EDTA-free protease inhibitor cocktail) was prepared. Then, 300 µL of rabbit reticulocyte lysate was layered on top of the sucrose gradient and centrifuged at 288,000 × g (40,000 rpm) for 2 hours at 4°C. Following centrifugation, the gradient was fractionated from the bottom of the tube. Fractions corresponding to the 10%-50% sucrose layers were collected in 500 µL volumes, and their absorbance at 260 nm (A_260_) was measured. The ribosome-containing fractions were pooled, and an equal volume of low-salt buffer (20 mM Tris-HCl, 150 mM KCl, 10 mM MgCl_2_, 1 mM DTT) was added, followed by polyethylene glycol 8000 (PEG8000) to a final concentration of 7% (from a 50% w/v stock solution). Ribosomes were then pelleted by centrifugation at 355,040 × g for 1 hour at 4°C ^[6,7]^.

Preparation of ribosome-depleted rabbit reticulocyte lysate supernatant: To obtain the translation-competent supernatant devoid of ribosomes, rabbit reticulocyte lysate was subjected to ultracentrifugation at 180,000 × g for 2 hours at 4□^[9]^. The resulting supernatant was carefully collected.

*In vitro* translation reconstitution: For subsequent translation assays, the isolated ribosomes were combined with the ribosome-depleted supernatant, synthetic linear mRNA templates, and an amino acid mixture. The reaction was incubated at 30°C for 2 hours, after which the translation products were analyzed.

### *Drosophila* Translation System

#### Preparation of *Drosophila* ovarian extract

The *Drosophila* ovarian extract was prepared according to previously described methods with minor modifications ^[4,5]^. Briefly, young female flies were cultured at 25°C on a yeast-containing diet for 2 to 3 days. Ovaries were dissected in ice-cold PBS and collected. All subsequent steps were performed on ice or at 4°C. The collected ovaries were allowed to settle by gravity, and the packed volume was measured. The ovarian pellets were then washed twice with a 12-volume mixture of PBS and Extraction Buffer DEI (10 mM Hepes pH 7.4, 5 mM DTT, supplemented with 1X COMPLETE-Protease Inhibitors Cocktail EDTA-free from Roche) at a 1:1 ratio. This was followed by two quick washes with a 12-volume of DEI buffer alone. The washed ovaries were transferred to a Dounce homogenizer (Kimble, 885300-0002). After removal of excess buffer, the ovaries were directly homogenized using the tight pestle. The homogenate was transferred to a microcentrifuge tube and centrifuged at 14,000 rpm for 10 minutes at 4°C. The resulting supernatant was carefully collected, representing the ovarian extract ready for subsequent *in vitro* translation assays. The extract could be aliquoted and stored at -80°C or used immediately.

#### *In vitro* translation

In a 25 µL reaction mixture, 1 µg of template circular RNA was translated in a system containing 60 µM amino acid mixture, 16.8 mM phosphocreatine, 80 ng/mL creatine kinase, 24 mM HEPES pH 7.4, 0.6 mM magnesium acetate, 60 mM potassium acetate, 0.1 mM spermidine, 1.2 mM DTT, 100 µg/mL bovine liver tRNA, and 40% ovarian extract. The reaction was incubated at 25°C for 120 minutes. Luciferase production was detected using a chemiluminescence assay (Beyotime, RG072S). For detection, Gaussia-Lumi™ substrate and buffer were mixed at a 1:100 ratio to prepare the working solution. A total of 10μL of translation product was mixed with 100μL of detection reagent (final volume: 110μL), incubated for 5 minutes at room temperature in the dark, and chemiluminescence was recorded using the GloMax Multi Jr detection system (Promega, E6070) with an integration time of 10 seconds per well.

## Acknowledgement

The support from National Natural Science Foundation of China Grant #32170658 (Y.Y.), the National Key Research & Developmental Program of China (2024YFF1206802), National Natural Science Foundation of China Grant #32200959 (L.S.), National Natural Science Foundation of China Grant #32100586 (F.C.), Fujian Natural Science Foundation of China Grant #2023J01412 (L.S.) are appreciated. We are grateful to Drs. Ruiwen Wang, Guixuan Zhou, Wenfeng Chen, Xiaozhen He, Xiang Lin for helpful discussions.

## Contributions

Yufeng Yang initiated the project. Yufeng Yang and Xiangyou Pi designed the experimental plan. Xiangyou Pi, Peng Wang, Bin Lin, Tianliang Liu, Yanchao Yang, Zhe Lin, Xiaoru Zhang, Zida Lin, Ling Sun, Fei Chen, Xi Zhang performed the experiments and data analysis. The manuscript was written by Yufeng Yang, Xiangyou Pi and Zhiliang Ji and contributed by all co-authors. Management and oversight of the project was performed by Yufeng Yang and Zhiliang Ji.

## Competing interests

The authors declare no competing interests.

## Supplementary Figures

**Figure S1.**
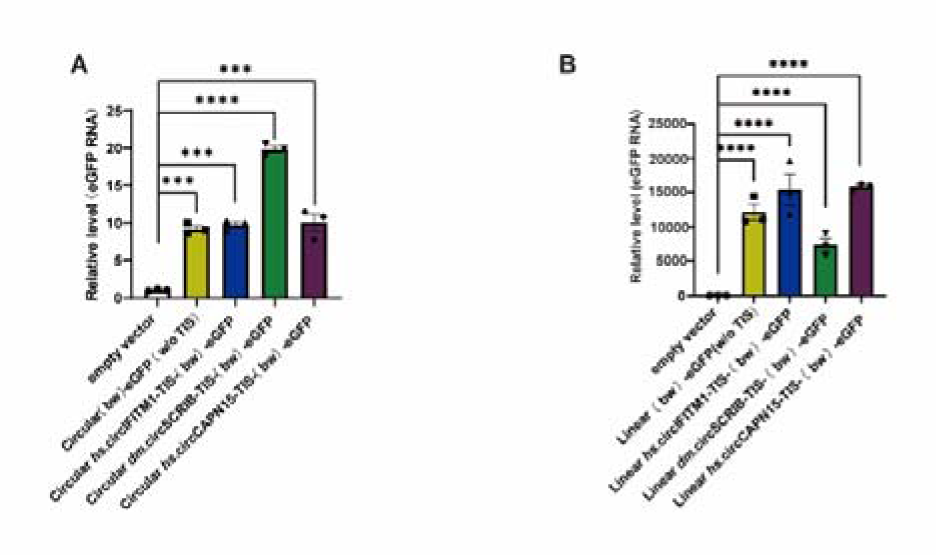
Backward eGFP quantification through RT-PCR. **(A)** RT-qPCR quantification of backward eGFP expression in HEK-293T cells transfected with empty vector, circular backward eGFP without TIS ((bw)-eGFP (w/o TIS)), *hs*.circIFITM1-TIS-(bw)-eGFP, *hs*.circCAPN15-TIS-(bw)-eGFP, and *hs*.circSCRIB-TIS-(bw)-eGFP (n=3). Ho-β-actin was used as the internal reference gene. Primers spanning the back-splice junction (BSJ) were used to specifically detect circular RNA. All RNA samples were treated with RNase R to enrich for circular species. **(B)** RT-qPCR was used to quantify the expression of backward eGFP in HEK-293T cells transfected with empty vector, linear TIS free backward eGFP (bw - eGFP (w/o TIS)), linear *hs*.circIFITM1-TIS - (bw) - eGFP, linear *hs.*circCAPN15-TIS - (bw) - eGFP, and linear *hs*.circSCRIB-TIS - (bw) - eGFP (n=3). *Ho*-β-actin is used as an internal reference gene.

**Figure S2.**
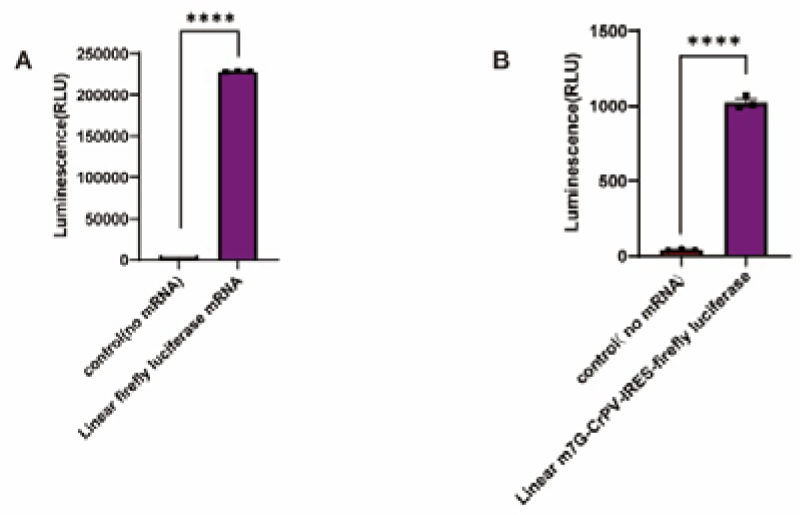
Positive controls in the cell-free *in vitro* translation systems. **(A)** Linear firefly luciferase RNA provided by the vendor served as the positive control for the the rabbit reticulocyte lysate assay (Fig. 3B). **(B)** Synthetic linear template m7G-CrPV-IRES-firefly luciferase mRNA served as the positive control for the reconstituted ribosome complex assay (Fig. 3D). The chemiluminescent intensity was then measured using the GloMax Multi Jr detection system following the reaction with firefly luciferase substrate (D-Luciferin). Statistical comparisons between groups were performed using Mann-Whitney U in GraphPad Prism (version 9.2.0). Statistical significance: ns: not significant, “*”: p <0.05, “**”: p <0.01, “***”: p <0.001, “****”: p <0.0001.

## Notes

### Competing Interest Statement

The authors have declared no competing interest.

## Reference

[1] Lin Z, Pi XY, Lv ZW, et al. 3’ to 5’ Translation of Circular RNAs? bioRxiv. 2025:2025.2012.2008.692888.

[2] McClain WH. Surprising contribution to aminoacylation and translation of non-Watson-Crick pairs in tRNA. Proc Natl Acad Sci U S A. 2006;103(12):4570–4575.

[3] Sakai Y, Kimura S, Suzuki T. Dual pathways of tRNA hydroxylation ensure efficient translation by expanding decoding capability. Nature Communications. 2019;10(1):2858.

[4] Gebauer F, Corona DF, Preiss T, Becker PB, Hentze MW. Translational control of dosage compensation in *Drosophila* by Sex-lethal: cooperative silencing via the 5’ and 3’ UTRs of msl-2 mRNA is independent of the poly(A) tail. EMBO J. 1999;18(21):6146–6154.

[5] Castagnetti S, Hentze MW, Ephrussi A, Gebauer F. Control of oskar mRNA translation by Bruno in a novel cell-free system from *Drosophila* ovaries. Development. 2000;127(5):1063–1068.

[6] Milicevic N, Jenner L, Myasnikov A, Yusupov M, Yusupova G. mRNA reading frame maintenance during eukaryotic ribosome translocation. Nature. 2024;625(7994):393–400.

[7] Khatter H, Myasnikov AG, Mastio L, et al. Purification, characterization and crystallization of the human 80S ribosome. Nucleic Acids Res. 2014;42(6):e49.

[8] Jia L, Nguyen TT, Uematsu S, et al. Programmable initiation of mRNA translation by trans-RNA. Nat Biotechnol. 2025.

[9] Trainor BM, Pestov DG, Shcherbik N. Development, validation, and application of the ribosome separation and reconstitution system for protein translation *in vitro*. RNA. 2021;27(12):1602–1616.

